# Proteasome-associated ubiquitin ligase relays target plant hormone-specific transcriptional activators

**DOI:** 10.1101/2021.10.04.462757

**Authors:** Zhishuo Wang, Beatriz Orosa-Puente, Mika Nomoto, Heather Grey, Thomas Potuschak, Takakazu Matsuura, Izumi C. Mori, Yasuomi Tada, Pascal Genschik, Steven H. Spoel

## Abstract

The ubiquitin-proteasome system is vital to hormone-mediated developmental and stress responses in plants. Ubiquitin ligases target hormone-specific transcriptional activators (TAs) for degradation, but how TAs are processed by proteasomes remains unknown. We report that in Arabidopsis the salicylic acid-and ethylene-responsive TAs, NPR1 and EIN3, are relayed from pathway-specific ubiquitin ligases to proteasome-associated HECT-type UPL3/4 ligases. Activity and stability of NPR1 was regulated by sequential action of three ubiquitin ligases, including UPL3/4, while proteasome processing of EIN3 required physical handover between ethylene-responsive SCF^EBF2^ and UPL3/4 ligases. Consequently, UPL3/4 controlled extensive hormone-induced developmental and stress-responsive transcriptional programmes. Thus, our findings identify unknown ubiquitin ligase relays that terminate with proteasome-associated HECT-type ligases, which may be a universal mechanism for processive degradation of proteasome-targeted TAs and other substrates.

**One-Sentence Summary:** Transcriptional activators are targeted by proteasomal ubiquitin ligase relays that control their activity and stability.

## Main Text

The ubiquitin-proteasome system plays vital roles in regulating cellular homeostasis and responses to the environment in eukaryotes. In plants developmental and stress response hormones extensively utilize the ubiquitin-proteasome system to precisely coordinate transcriptional programs (*1*). Several plant hormones have been shown to act as molecular glue between ubiquitin ligases and their substrates (*2-4*). This leads to substrate modification by a chain of the small 8 kDa protein ubiquitin that targets substates to the proteasome for degradation (*5*). Hormone-induced degradation of corepressors releases the activity of transcriptional activators (TAs), thereby triggering genome-wide transcriptional changes. Additionally, hormones control the activities of ubiquitin ligases that directly target TAs to regulate their stability (*6*). For example, EIN3 is a master TA of the developmental and stress hormone ethylene (*7*). In absence of ethylene, EIN3 is rapidly targeted to the proteasome by the modular Skp-Cullin-F-box (SCF) ubiquitin E3 ligase, SCF^EBF1/2^, in which EBF1/2 adaptors specifically recruits EIN3 for ubiquitination (*8-10*). Thus, when ethylene levels fall, SCF^EBF1/2^-effectively shuts down EIN3-induced transcriptional reprogramming. By contrast, the plant immune hormone salicylic acid (SA) stimulates the step-wise ubiquitination of NPR1, a master TA of hundreds of immune genes and promoter of cell survival (*11-13*). Initial SA-induced ubiquitination of NPR1 by a Cullin-RING Ligase 3 (CRL3) activates NPR1, while subsequent ubiquitin chain elongation by Ubiquitin conjugation factor E4 (UBE4) ligase deactivates NPR1 and targets it for degradation (*14*). In addition, SCF^HOS15^ ligase targets NPR1 for degradation to limit and prevent untimely activation of immune genes (*15*). Hence, progressive ubiquitination and subsequent turnover of NPR1 are critical steps in SA-induced immune gene activation.

While the steps leading up to degradation of plant hormone-specific TAs are increasingly well understood, how TAs are shuttled to the proteasome and how the proteasome affects their intrinsic transcriptional activities remains largely unknown. Intriguingly, the proteasome itself harbors ubiquitin ligase activity that is thought to be important for promoting proteasome processivity (*16-18*). Proteasome-associated ubiquitin ligase activity is conferred by HECT-type ubiquitin ligases that directly interact with the 19S proteasome subcomplex. Recently, we reported that a member of the Arabidopsis HECT-type family of Ubiquitin Protein Ligases (UPL) not only interacts with the proteasome but in yeast two-hybrid assays also physically associated with hormone-responsive ubiquitin ligases (*19*). This suggest that UPLs may play an important and yet unrecognized role in proteasome-mediated plant hormone signaling.

Genetic experiments revealed that UPL1, 3 and 5 are important for trichome development and SA-induced immunity (*19, 20*). Therefore, we first assessed the activities of their respective HECT domains in assembling ubiquitin chains. In presence of the full ubiquitination machinery, HECT domains from all three UPLs successfully formed ubiquitin conjugates, while mutation of the active site cysteine partly compromised their activities (fig. S1A)(*21*). Moreover, like we reported previously for UPL3 (*19*), both the N-terminus of UPL1 and full-length UPL5 co-immunoprecipitated with the proteasome *in vivo* (fig. S1B). Therefore, we assessed if association of all three UPLs endows the proteasome with ubiquitin ligase activity. However, only proteasomes from immune-induced *upl3* knockout mutants displayed a substantial reduction in proteasome-associated ubiquitin ligase activity (fig. S1C), indicating that at least *in vitro*, UPL3 is the primary active ligase. Nonetheless, *upl1, upl3* and *upl5* mutants all displayed decreased cellular levels of ubiquitin conjugates as well as polyubiquitination of the model substrate RPN10 (fig. S1D), suggesting these UPLs broadly catalyze polyubiquitination of numerous cellular proteins.

Lack of UPL3 activity is associated with failure to reprogram the transcriptome upon activation of immunity (*19*). Similarly, *upl1* and *upl5* mutants were partially defective in SA-induced marker gene expression and transcriptome reprogramming (Fig 1, A and B, and fig S2A). The majority of SA-induced, UPL-regulated genes were dependent on SA-responsive NPR1 coactivator (fig. S2B)(*19*), suggesting that UPLs may regulate the stability of NPR1. Although NPR1 transcript levels were unaffected, pathogen-and SA-induced accumulation of endogenous NPR1 protein was compromised in all three *upl* mutants (Fig. 1, C and D, and fig. S2C). Reduced accumulation of NPR1 protein was likely due to decreased SA levels in *upl* mutants (fig. S2D), as SA is required for NPR1 protein homeostasis (*22*), or due to changes in mRNA translation. To circumvent this indirect effect and explore if UPLs directly affect NPR1 protein stability and activity, we constitutively expressed *NPR1-GFP* (without UTRs) in *upl* mutants (fig. S2, E and F). Although this restored SA-induced expression of *PR1* (direct NPR1 target gene) in *upl1* and *upl5* mutants, higher levels of NPR1-GFP protein were required compared to the wild-type (WT) background (Fig. 1E, and fig. S2F). More strikingly, NPR1-GFP largely failed to induce *PR1* gene expression in *upl3* mutants (Fig. 1E). These data strongly imply that UPL ligases, and in particular UPL3, regulate NPR1’s TA activity.

**Fig 1.**
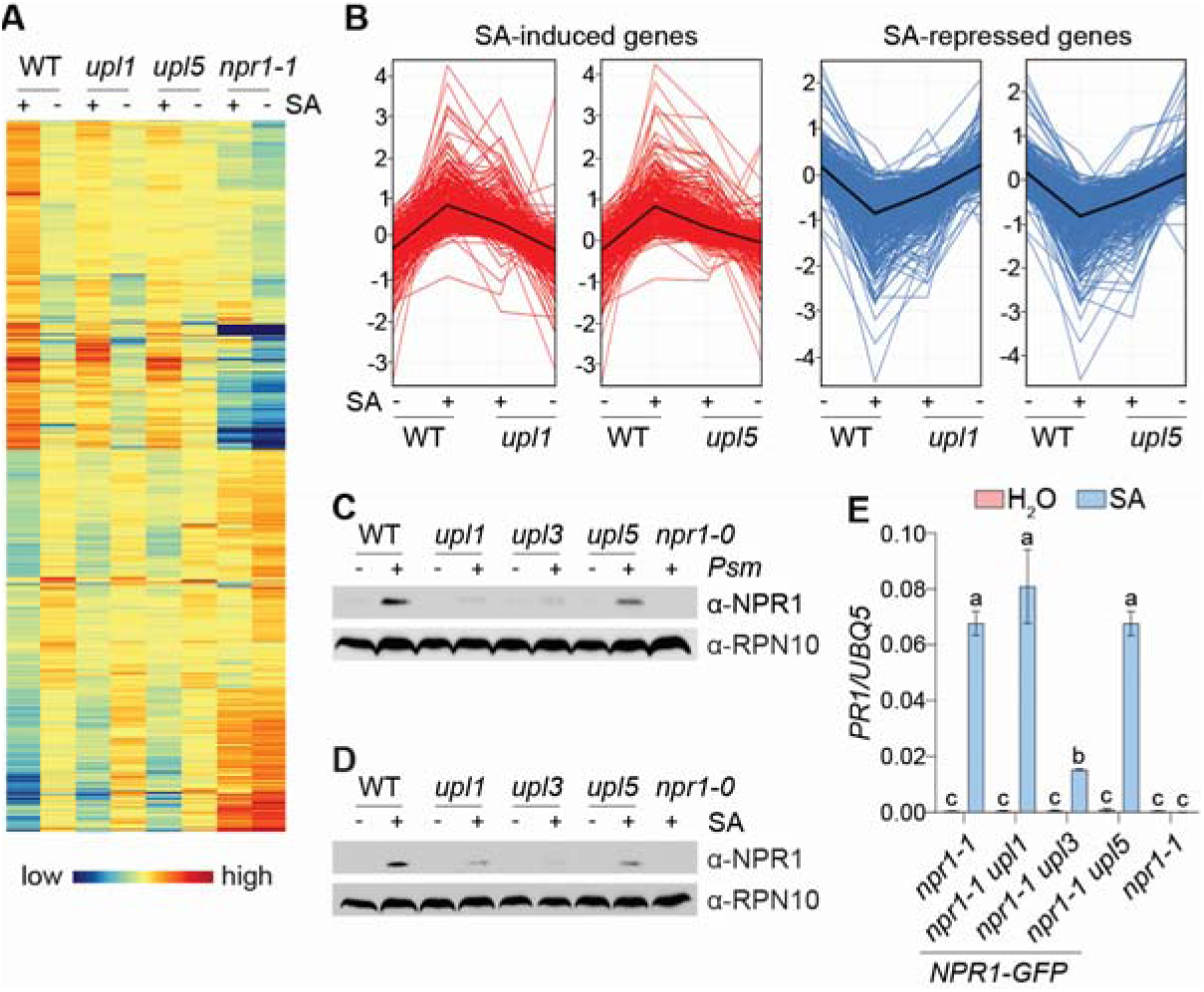
UPL ligases are required for SA- and NPR1-dependent transcriptional reprogramming. **(A and B)** Mutant *upl* plants are impaired in SA-dependent transcriptional reprogramming. Adult plants were treated with 0.5 mM SA or H_2_O for 24 hours, mRNA was extracted and analysed by RNA-Seq. Genes with fold change of ≥ 1.5 (Benjamini Hochberg FDR, 2-way ANOVA *p* ≤ 0.05, n = 3) in WT plants in response to SA are shown as a heat map (A) and profile plot (B). **(C)** Pathogen-induced accumulation of endogenous NPR1 protein is controlled by UPLs. Indicated genotypes were inoculated with or without 10^6^ cfu/ml *Pseudomonas syringae* pv. *maculicola* ES4326 (*Psm*) and endogenous NPR1 protein detected by immunoblotting, while RPN10 was used as a loading control. **(D)** SA induced accumulation of endogenous NPR1 protein is controlled by UPLs. Indicated genotypes were treated with 0.5 mM SA or H_2_O for 24 hours and proteins detected as in (C). **(E)** NPR1-mediated *PR1* gene expression is impaired in *upl3* plants. Seedlings constitutively expressing *NPR1-GFP* in indicated genetic backgrounds were treated with 0.5 mM SA or H_2_O for 6 hours. *PR-1* gene expression was normalised to *UBQ5*. Data represent mean ± SD (Tukey HSD ANOVA test; α = 0.05, n = 3).

To explore this possibility, we assessed first if UPLs interact with NPR1 *in vivo*. While we were unable to express UPL1, both epitope-tagged UPL3 and UPL5 co-immunoprecipitated with NPR1-GFP (Fig. 2A). Moreover, the levels of SA-induced polyubiquitinated NPR1-GFP were markedly reduced in *upl* mutants (Fig 2B). Treatment with the protein synthesis inhibitor, cycloheximide, demonstrated that while NPR1-GFP was rapidly degraded when expressed in the WT, it was significantly more stable in *upl* mutants (Fig. 2, C and D). Together, these findings show UPL ligases polyubiquitinate SA-induced NPR1 and promote its degradation by the proteasome. Given that UPLs are associated with the proteasome, we reasoned that they might function sequentially after CRL3 and UBE4 ligases to modify NPR1 and promote its degradation. We previously reported that in contrast to *upl* mutants, mutant *ube4* plants accumulate highly active NPR1 that is modified by short ubiquitin chains (*14*). Unfortunately, we were unable to obtain homozygous *upl3 ube4* double mutants, suggesting this combination was lethal. However, in agreement with a ubiquitin ligase relay consisting of CRL3, UBE4 and ending with UPL3, heterozygous knockout of *UBE4* in *upl3* mutants largely restored NPR1’s ability to activate *PR1* and *PR2* gene expression (Fig. 2, E and F, and fig. S2G).

**Fig 2.**
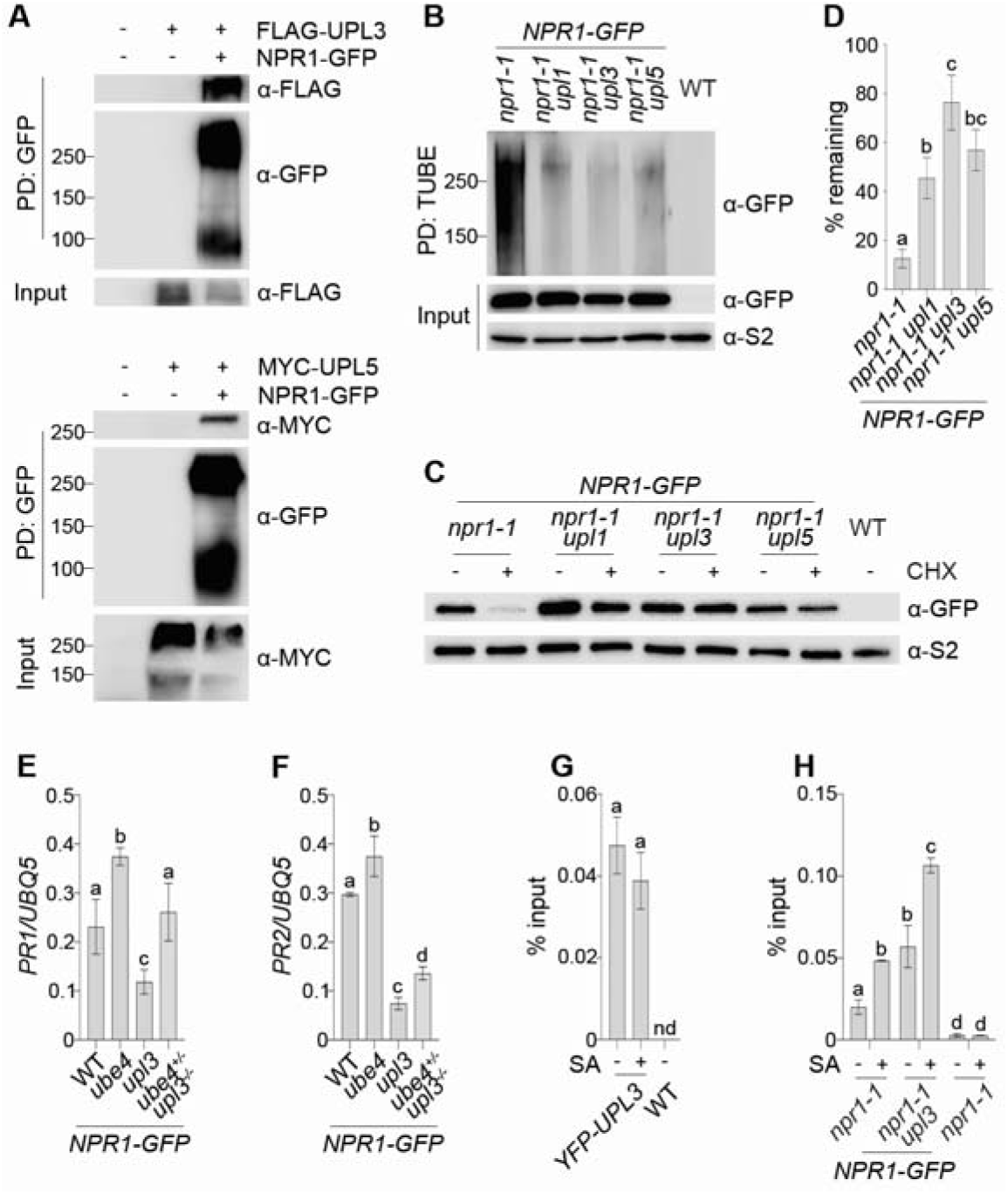
UPL ligases polyubiquitinate SA-induced NPR1 and promote its degradation by the proteasome. (**A**) UPLs physically interact with NPR1. NPR1-GFP was transiently expressed with FLAG-UPL3 or MYC-UPL5 in *Nicotiana benthamiana*. Protein complexes were pulled down with GFP-Trap agarose and analysed by immunoblotting against GFP, FLAG and MYC. (**B**) UPLs polyubiquitinate NPR1. Indicated genotypes expressing NPR1-GFP were treated with 0.5 mM SA and 100 μM MG132 for 6 h. Ubiquitinated proteins were pulled down with GST-TUBE and analysed by immunoblotting against GFP and S2 (regulatory non-ATPase subunit RPN1 used as loading control). (**C**) NPR1 is stabilised in *upl* mutants. Indicated genotypes expressing NPR1-GFP were treated with 100 μM CHX or DMSO vehicle for 2 h. Protein was analysed by immunoblotting against GFP and S2 (loading control). (**D**) As in (C) but NPR1-GFP protein remaining after CHX treatments were quantified relative to DMSO treatments (Tukey HSD ANOVA test; α = 0.05, n = 3). (**E** and **F**) Mutation of *UBE4* in *upl3* background restores expression of *PR* genes. Indicated genotypes expressing NPR1-GFP were treated with 0.5 mM SA for 24 h. Expression of *PR* genes was normalised to *UBQ5*. Data represent mean ± SD (Tukey HSD ANOVA test; α = 0.05, n = 3). (**G**) YFP-UPL3 localises to the *PR-1* promoter. Adult *35S:YFP-UPL3 (upl3)* plants were treated with 0.5 mM SA or H_2_O for 24 hours before assessing YFP-UPL3 binding to the *as-1* motif of the *PR-1* promoter. Data represent mean ± SD (Tukey HSD ANOVA test; α = 0.05, n = 3); nd, not detected. (**H**) NPR1-GFP accumulates at the *PR1* promoter of *upl3* mutants. As in (G) but binding of NPR1-GFP to *PR-1* promoter was analysed.

Our data show that although *upl3* mutants fail to degrade NPR1, they are compromised in SA-induced expression of *PR1*. To understand how UPL3-mediated ubiquitination of NPR1 may influence its TA activity, we assessed chromatin association of both UPL3 and NPR1. UPL3 was constitutively associated with the *PR1* promoter independent of SA treatment (Fig. 2G). By contrast, when expressed in the *npr1-1* mutant background, NPR1-GFP was recruited to the *PR1* promoter only in response to SA treatment. Unexpectedly, however, NPR1-GFP accumulated to much higher levels at the *PR1* promoter of *upl3 npr1-1* double mutants both before and particularly after SA treatment (Fig 2H). Collectively these findings show that proteasome-associated UPL3 is the last in a relay of three ubiquitin ligases that polyubiquitinate NPR1 and ensures transcriptionally inactive NPR1 is cleared from target gene promoters by the proteasome.

We then asked if it is a general phenomenon that unstable TAs are subjected to ubiquitin ligase relays that end in their ubiquitination by proteasome-associated UPLs. Previous studies suggest that some ubiquitin ligases, including hormone-responsive ones, can associate with the proteasome (*23-26*). Thus, it is plausible that these ubiquitin ligases physically relay substrates to UPLs and the proteasome. In agreement with this we previously found by yeast two-hybrid that the UPL3 N-terminus interacts with the F-box protein EBF2, the substrate adaptor of an ethylene-responsive SCF^EBF1/2^ ligase that targets the transcriptional activator EIN3 for degradation (*19*). First, we verified by coimmunoprecipitation that physical interaction between full-length UPL3 and EBF2 indeed takes place in plants (Fig. 3A). Moreover, we found that along with a proteasome regulatory subunit, endogenous EIN3 also co-immunoprecipitated with UPL3 (Fig 3B). To investigate if SCF^EBF1/2^ directly hands over EIN3 to UPL3 for further ubiquitination, we compared interaction between HA-tagged EIN3 and YFP-tagged UPL3 in presence or absence of EBF1 and EBF2. Strikingly, interaction between HA-EIN3 and YFP-UPL3 was completely dependent on EBF1/2 (Fig. 3C), suggesting SCF^EBF1/2^ hands over its cargo to UPL3 in a previously unknown physical relay.

**Fig 3.**
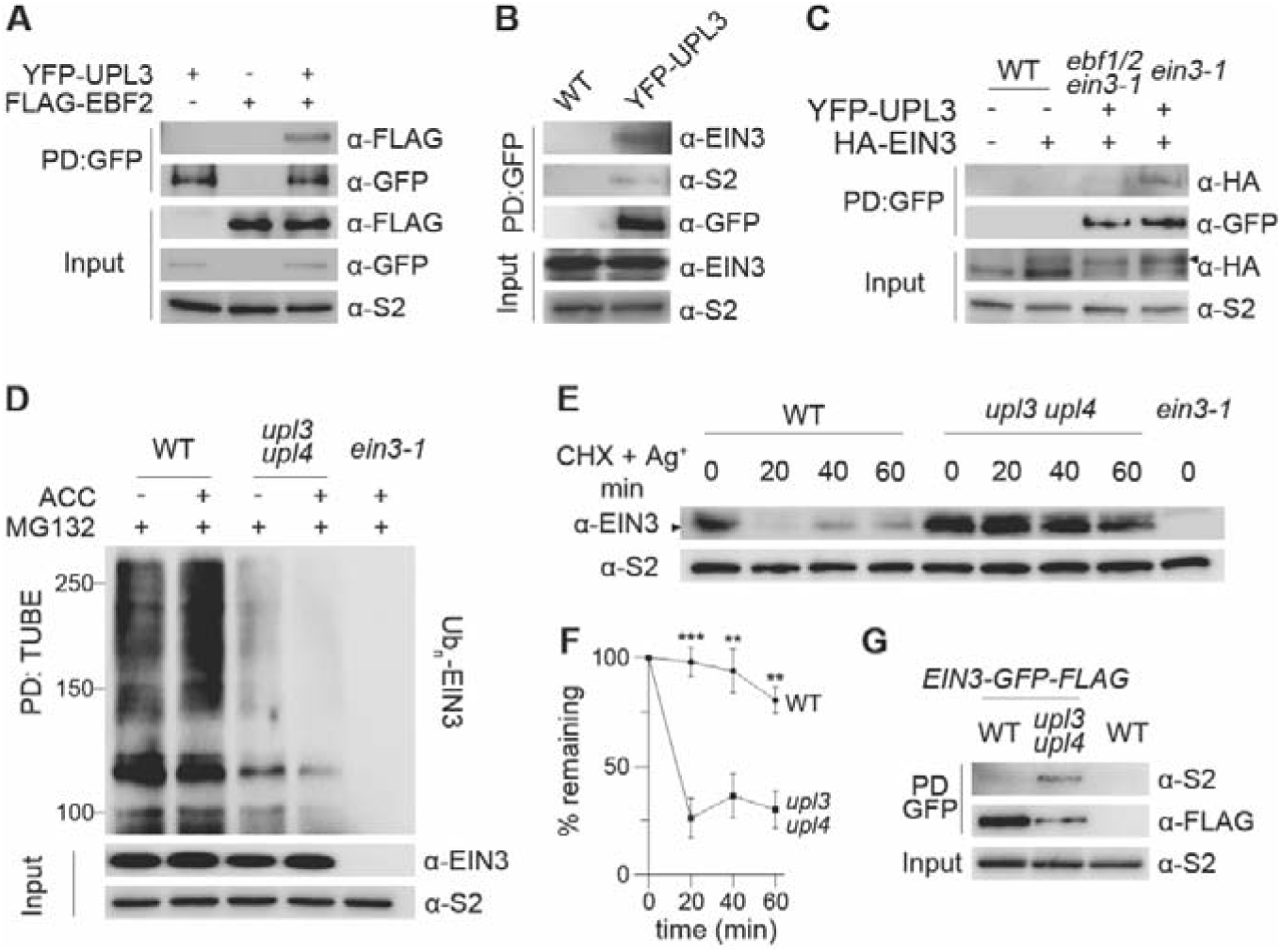
An SCF^EBF1/2^–UPL3/4 ubiquitin ligase relay is required for proteasomal processing of EIN3. (**A**) UPL3 interacts with EBF2. YFP-UPL3 was purified from *35S:YFP-UPL3* plants with GFP-Trap and incubated with *in vitro* synthesized FLAG-EBF2. Immunoprecipitated proteins were analyzed by immunoblotting against GFP and FLAG. (**B**) UPL3 interacts with EIN3. Proteins from *35S:YFP-UPL3 (upl3)* seedlings were pulled down with GFP-Trap and analyzed by immunoblotting against EIN3, GFP and S2 (loading control). (**C**) UPL3-EIN3 interaction is dependent on EBF1/2. *35S:YFP-UPL3 (ein3-1)* and *35S:YFP-UPL3 (ebf1 ebf2 ein3-1)* protoplasts were transformed with *35S:HA-EIN3*. Proteins were pulled down with GFP-Trap and analyzed by immunoblotting against HA, GFP and S2 (loading control). (**D**) UPL3/4 polyubiquitinate EIN3. Seedlings were treated with 100 μM MG132 and 50 μM ACC for 3 h. Ubiquitinated proteins were pulled down with His-TUBE and analysed by immunoblotting against EIN3 and S2 (loading control). (**E**) EIN3 is stabilized in *upl3 upl4* mutants. Seedlings were submerged in 50 μM ACC for 3 h and then transferred to a combination of 100 μM CHX and 100 μM AgNO3 for the indicated times. Proteins were analyzed by immunoblotting against EIN3 and S2 (loading control). (**G**) Proteasomal degradation of EIN3 stalls at *upl3 upl4* proteasomes. Proteins from plants expressing *pEIN3:EIN3-eGFP-3xFLAG* in WT and *upl3 upl4* backgrounds were pulled down with GFP-Trap and analyzed by immunoblotting against S2 and FLAG.

So why are TAs relayed from pathway-specific ubiquitin ligases to proteasome-associated UPLs? It is plausible that UPLs add or remodel ubiquitin chains on TAs to ensure they retain high affinity while being degraded. In agreement, regardless of treatment with the ethylene precursor 1-aminocyclopropane 1-carboxylic acid (ACC), polyubiquitination of endogenous EIN3 was markedly reduced when proteasome activity was blocked in a double knockout mutant of both *UPL3* and its closest homologue *UPL4* (Fig. 3D). We then assessed if this led to changes in EIN3 stability in *upl3 upl4* mutants. Seedlings were first treated with ACC to allow accumulation of EIN3, after which destabilization of EIN3 was triggered by treatment with silver ions, a potent inhibitor of ethylene action (*27*), as well as the protein synthesis inhibitor CHX. While EIN3 was degraded within minutes of treatment in the WT, it was much more stable in *upl3 upl4* mutants (Fig. 3, E and F). To investigate the effect of UPL3/4-mediated EIN3 polyubiquitination on the proteasome, we expressed epitope-tagged EIN3 in WT and *upl3 upl4* plants (fig S3A), and assessed its association with the proteasome via the regulatory subunit RPN1 (S2) that is located at the base of the 19S particle (*28*). Due to continuous EIN3 degradation, interaction between EIN3 and the proteasome was only barely detectable in WT plants (Fig. 3G). By contrast, EIN3 accumulated at proteasomes in *upl3 upl4* mutants, indicating its proteasomal degradation was stalled (Fig. 3G). From these experiments we draw two conclusions. Firstly, while SCF^EBF1/2^ physically relays EIN3 to UPL3 (Fig. 3C), polyubiquitinated EIN3 can still recruit or be recruited to proteasomes in absence of UPL3/4 (Fig. 3G), suggesting interaction with SCF^EBF1/2^ may activate UPL3 to engage with EIN3. And secondly, relay of EIN3 from SCF^EBF1/2^ to UPL3/4 results in ‘eleventh hour’ polyubiquitination, which is necessary for its processive degradation by the proteasome.

The proteasome plays an important role in limiting ethylene responses by maintaining low steady-state levels of EIN3 (*8-10*). As expected, we found that UPL3/4 contribute to this process, as *upl3 upl4* mutants accumulated high levels of endogenous EIN3 even in absence of ACC-induced ethylene signaling (Fig. 4A and fig. S3B). Compared to WT, the basal and ACC-induced expression of EIN3 target genes was consequently also enhanced in *upl3 upl4* mutants (Fig 4B and fig. S3, C and D). We then sought to uncover the developmental effect of UPL3/4-mediated regulation of EIN3 by assessing the ‘triple response’ of etiolated seedlings (*29*). In presence of active ethylene signaling, dark-grown seedlings display a short, thickened root and hypocotyl with an exaggerated apical hook. Similar to *ebf1 ebf2* mutants that fail to degrade EIN3, *upl3 upl4* mutants displayed a phenotype consistent with constitutive ethylene signaling (fig. S3, E to G). To determine if this phenotype was dependent on EIN3 we generated *upl3 upl4 ein3* triple mutants. The enhanced expression of EIN3 marker genes observed in *upl3 upl4* double mutants was largely lost in this triple mutant (Fig. 4C and fig. S4, A and B). A similar picture was observed across the entire ACC-responsive transcriptome with mutation of EIN3 dampening transcriptional reprogramming caused by knockout of *UPL3* and *UPL4* (Fig. 4, D to F). Consequently, the constitutive ethylene response phenotype of *upl3 upl4* plants was partially lost by mutation of EIN3 (fig. S4, C to E). Residual ethylene signaling was likely due to a notable number of ACC-responsive genes that are independent of EIN3 but regulated by UPL3/4 (Fig. 4E). This suggests that UPL3/4 may also target previously described EIN3-like (EIL) TAs (*30*).

**Fig 4.**
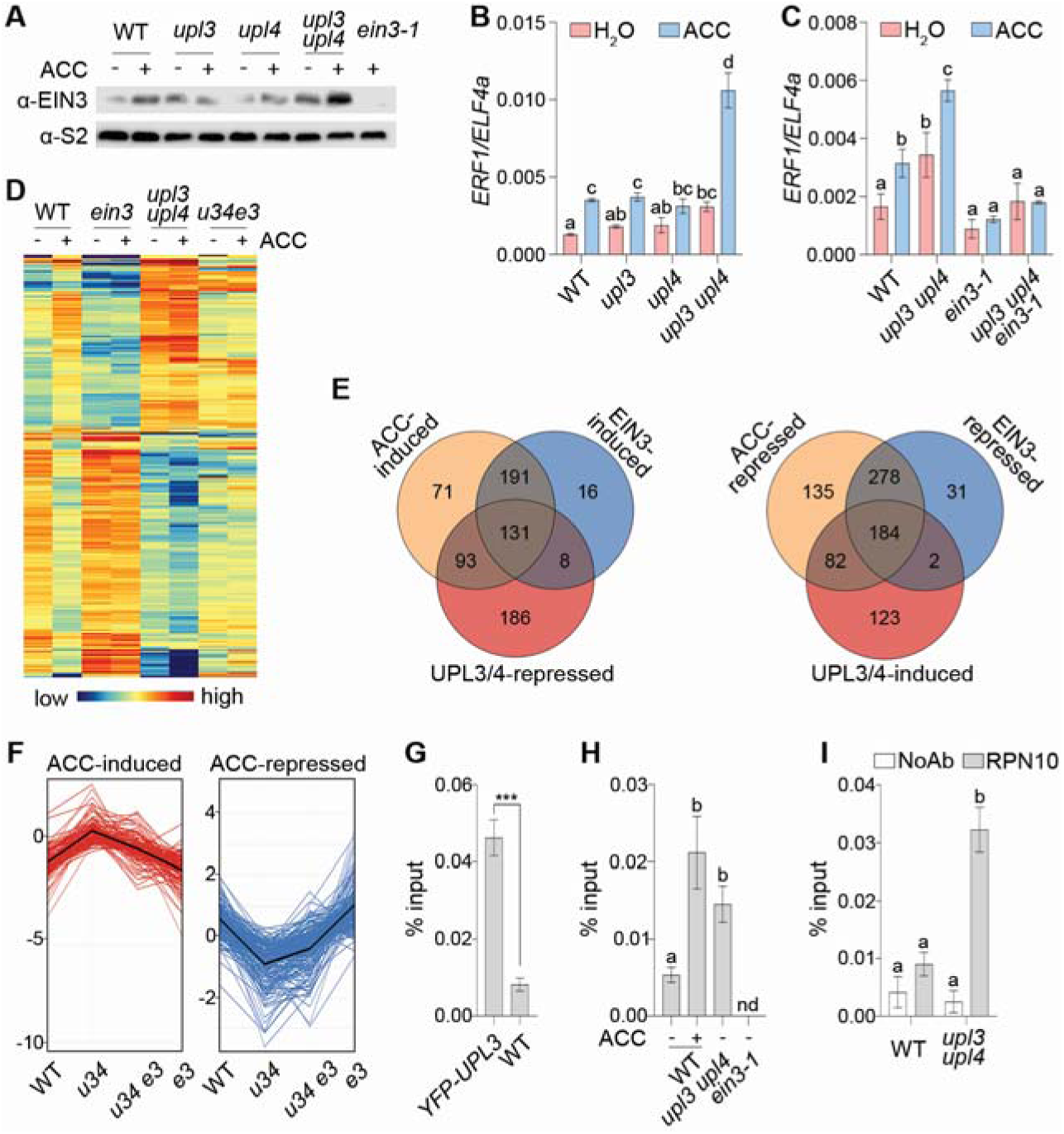
UPL3/4 are required for EIN3-mediated transcriptional reprogramming. (**A**) Mutant *upl3 upl4* plants accumulate EIN3. Seedlings were treated with 50 μM ACC for 3 hours and proteins analyzed by immunoblotting against EIN3 and S2. (**B** and **C**) Mutant *upl3 upl4* plants exhibit enhanced expression of an EIN3 target gene. Seedlings were treated as in (A) and levels of *ERF1* gene expression determined and normalised to *ELF4a*. Data represent mean ± SD (Tukey HSD ANOVA test; α = 0.05, n = 3). (**D**) Mutant *upl3 upl4* plants display constitutive ET-responsive gene expression. Seedlings were treated with 50 μM ACC or H_2_O for 3 h and mRNA analysed by RNA-Seq. ACC-responsive genes (fold change ≥ 1.5, Benjamini Hochberg FDR, 2-way ANOVA *p* ≤ 0.05, n = 3) that were regulated by both UPL3/4 and EIN3 are shown as a heat map (D) and profile plot (F). Venn diagrams (**E**) illustrate overlaps between ACC-regulated genes, EIN3-regulated genes, and UPL3/4-regulated genes. (**G**) UPL3 localises to the ET-responsive *ERF1* promoter. *35S:YFP-UPL3 (upl3)* plants were analysed by ChIP with a GFP antibody Data represent mean ± SD, asterisks indicate statistically significant differences (two-tailed t-test, ***p ≤ 0.001, n = 3). (**H**) EIN3 accumulates at the *ERF1* promoter of *upl3 upl4* mutants. Seedlings were treated with 50 μM ACC or H2O for 3 h before assessing EIN3 binding to the *ERF1* promoter. Letters indicate statistical differences (Tukey HSD ANOVA test; α = 0.05, n = 3); nd, not detected. (**I**) Proteasomes accumulate at the *ERF1* promoter of *upl3 upl4* mutants. Seedlings were analysed by ChIP with an RPN10 proteasome subunit antibody. Data were analysed as in (H). NoAb, no antibody negative control.

Finally, we explored if UPL3/4 and the proteasome directly control ethylene-responsive transcription by regulating chromatin-associated EIN3. Indeed, transgenic YFP-UPL3 was localized to the promoter of *ERF1*, a direct target gene of EIN3 (Fig. 4G). In WT plants ACC treatment induced recruitment of EIN3 to the *ERF1* promoter, while in *upl3 upl4* mutants EIN3 already accumulated at this promoter even in absence of ACC (Fig. 4H). Thus, UPL3/4 limit accumulation of EIN3 at target genes, thereby avoiding their untimely activation. Importantly, we also found the proteasome was highly enriched at the *ERF1* promoter of *upl3 upl4* mutants (Fig. 4I), suggesting that stalling of EIN3 degradation traps the proteasome at ethylene-responsive genes.

Taken together we have uncovered a novel relay mechanism by which plant hormone-specific TAs are transmitted between different ubiquitin ligases to control their transcriptional activities. Relays terminate at the proteasome where ‘eleventh hour’ polyubiquitination by HECT-type ligases ensures processive TA degradation. Our data suggest that in at least two cases, TAs are physically handed over from pathway-specific ubiquitin ligases to proteasome-associated HECT-type ligases. Consequently, proteasome-associated HECT-type ligases play an indispensable role in plant hormone-induced transcriptional reprogramming. As HECT-type ligases are bound to proteasomes in a variety of different eukaryotes (*16, 19, 31*), ubiquitin ligase relays may be a universal mechanism for proteasome-mediated substrate degradation.

## Acknowledgments

The authors thank Dr. Michael Skelly for critically reading the manuscript.

## Funding

European Research Council (ERC) under the European Union’s Horizon 2020 research and innovation program, grant agreement No. 678511 (S.H.S.)

Biotechnology and Biological Sciences Research Council (BBSRC) grant BB/S016767/1 (S.H.S.)

Darwin Trust PhD studentship (Z.W.)

Royal Society International Exchanges grant IEC\R3\170118 (S.H.S, Y.T.)

SPS Grant-in-Aid for Scientific Research (B) grant 16H05065 (Y.T.)

Agence Nationale de la Recherche (ANR) grant ANR-10-IDEX-0002 and IMCBio ANR-17-EURE-0023 (T.P., P.G.)

## Author contributions

ZW, BO, and SHS designed the research. ZW and BO performed experiments. TP and PG provided crucial research materials. MN and TM performed hormone quantifications. HG, ICM, YT, PG and SHS performed project administration and acquired funding. The manuscript was written by ZW, BO and SHS, and further edited by TP and PG.

## Competing interests

Authors declare that they have no competing interests.

## Data and materials availability

All data are available in the main text or the supplementary materials except for RNA-Seq data, which have been deposited in Array Express at EMBL-EBI under accession codes E-MTAB-10963 and E-MTAB-10964. All materials used in the study are available upon request via materials transfer agreements (MTAs).

## Supplementary Materials

Materials and Methods

Supplementary Text

Figs. S1 to S4

Tables S1 to S2

References (*32*–*49*)

## Notes

https://www.ebi.ac.uk/arrayexpress/experiments/E-MTAB-10963

https://www.ebi.ac.uk/arrayexpress/experiments/E-MTAB-10964

### Competing Interest Statement

The authors have declared no competing interest.

