## Supplementary Materials for "Proteasome-associated ubiquitin ligase relays target plant hormone-specific transcriptional activators"

**This PDF file includes:**

Materials and Methods

Figs. S1 to S4

Tables S1 to S2

Materials and Methods

Plant materials, growth conditions, hormone treatments and phenotype analysis

All Arabidopsis plants used in this study are in the Columbia-0 (Col-0) background. The *upl1* (SALK_063972), *upl3* (SALK_035524), *upl4* (SALK_040984), *upl5* (SALK_116446), *npr1-0* (SALK_204100), *npr1-1*, *ein3-1*, *ebf1-1 ebf2-1* mutants, and *35S:YFP-UPL3* transgenic lines have been described previously (*32-35*). The HECT domains of UPL1 (amino acids 3238-3681), UPL3 (amino acids 1403-1888), UPL5 (amino acids 444-873) were cloned into the pENTR/D-TOPO vector (Invitrogen), and then were recombined into pEarleyGate 104 by LR reaction. The cysteine residues in the HECT domains of UPL1 (amino acid 3648), UPL3 (amino acid 1855), or UPL5 (amino acid 839) were mutated into serine residues by using a QuickChange Lightning Site-Directed Mutagenesis Kit (Agilent). To generate the *pCAB1:NPR1-GFP* construct, the coding sequence of *NPR1* fused with *GFP* was cloned into the pENTR/D-TOPO vector and subsequently recombined into the pCAB1:GW binary vector by LR reaction (*36*). The pEIN3:EIN3-eGFP-3xFlag construct was generated using the pART27 vector backbone (*37*). Briefly, the 35S promoter of the expression vector pPILY was replaced with the genomic sequence of EIN3 using XhoI-NcoI restriction digest (*38*). After insertion of eGFP and Flag coding sequences, the fragment of EIN3 genomic fusion with eGFP-3xFlag was inserted into binary vector pART27 using NotI restriction digest.

All plant materials and vectors used are listed in Table S1.

For experiments on adult plants, seeds were germinated under long day condition (16 h light, 8 h dark) at 65% humidity and 22°C with light intensity of 70–100 μmol m^-2^sec^-1^. For experiments on seedlings, seeds were washed in 100% ethanol for 5 min, followed by incubation in 10% bleach for 5 min, and then plated on Murashige and Skoog (MS) agar media. All plated seeds were kept at 4°C for 2-4 days before moving to the growth chamber.

For SA treatment, 4-week-old adult plants were sprayed with 0.5 mM SA (sodium salicylate, Sigma-Aldrich), while seedlings were immersed in 0.5 mM SA or H_2_O. For pathogen inoculation, *Psm* ES4326 was grown in LB media supplemented with 10 mM MgCl_2_. Cells were collected from the overnight cultures and diluted in 10 mM MgCl_2_ to appropriate concentration. Plants were infiltrated with a syringe through the abaxial leaf surface.

For treatment with 1-aminocyclopropanecarboxylic acid (ACC, Sigma-Aldrich), 10-day-old seedlings were treated with H_2_O or 50 μM ACC for 3 h. For analysing the triple response, seeds were germinated in the dark on MS media supplemented with or without 10 μM ACC. The hypocotyl and root lengths of 4-day-old seedlings were measured with Image J.

Gene expression measurement

Total RNA was extracted as described (*39*), cDNA was synthesized by using SuperScript™ II Reverse Transcriptase (Invitrogen) according to the manufacturers’ instructions. qPCR was performed by using PowerUp™ SYBR™ Green Master Mix (Applied Biosystems) on a StepOnePlus^TM^ Real-Time PCR system (Applied Biosystems). Primers used for qPCR were listed in Table S2.

For the RNA-Seq analysis, total RNA was further purified by using RNeasy Mini Kit (Qiagen). The RNA-Seq reads were aligned to the Arabidopsis thaliana TAIR10 genome using Bowtie. TopHat identified potential exon-exon splice junctions of the initial alignment. Strand NGS software in RNA-Seq workflow was used to quantify transcripts. Raw counts were normalised using DESeq with baseline transformation to the median of all samples. Data were then expressed as normalised signal values (i.e. log_2_[RPKM] where RPKM is read count per kilobase of exon model per million reads) for all statistical tests and plotting. RNA-Seq data have been deposited in Array Express at EMBL-EBI under accession codes E-MTAB-10963 and E-MTAB-10964.

Protein analysis

Liquid nitrogen frozen plant tissue was ground in protein extraction buffer (50 mM Tris-HCl (pH 7.5), 150 mM NaCl, 5 mM EDTA, 0.1% Triton X-100, 0.2% Nonidet P-40, 50 μg/ml TPCK, 50 μg/ml TLCK, 0.6 mM PMSF) unless otherwise stated. Protein extracts were incubated with 1× SDS sample buffer supplemented with 50 mM DTT at 80°C for 10 min, and then were separated by SDS-PAGE. All the antibodies used are listed in Table S1.

Endogenous NPR1 was detected by using anti-NPR1 antibody (Agrisera). For NPR1-GFP degradation assay, 2-week-old seedlings were treated with 100 μM CHX and samples collected 2 h after treatment. NPR1-GFP was detected using an anti-GFP antibody (Roche).

For analysing accumulation of EIN3, samples were ground in protein extraction buffer containing 62.5 mM Tris-Cl (pH 6.8), 3% SDS, 10% glycerol, 0.1% bromophenol blue, protease inhibitors (50 μg/mL N-p-Tosyl-L-phenylalanine chloromethyl ketone (TPCK), 50 μg/mL Nα-Tosyl-L-lysine chloromethyl ketone hydrochloride (TLCK), 0.6 mM phenylmethylsulfonyl fluoride (PMSF)), and 3% 2-mercaptoethanol. The mixture was incubated at 95°C for 5 min. For EIN3 degradation assay, 10-day-old seedlings were pre-treated with 50 μM ACC for 3 h, then were transferred into MS liquid media containing 100 μM CHX and 100 μM AgNO_3_, and samples were collected at indicated time points. Endogenous EIN3 was detected by using a previously described anti-EIN3 antibody (*40*).

GST-TUBE and His-TUBE pull-down of ubiquitinated substrates were performed as previously described (*34*). Total ubiquitination level was detected by using anti-ubiquitin antibody (anti-ubiquitinylated proteins clone FK2, Merck), while ubiquitinated NPR1-GFP and EIN3 were detected by immunoblotting with anti-GFP (Roche) and anti-EIN3 (*40*) antibodies, respectively.

In vitro ubiquitination assay

For purification of YFP-HECT, *Agrobacterium tumifaciens* carrying *35S:YFP-HECT* were collected from the overnight cultures and resuspended in infiltration buffer containing 10 mM MgCl_2_ and 10 μl/l 6-benzyladenine to OD_600_ = 0.5. *Nicotiana benthamiana* leaves were infiltrated with this Agrobacterium suspension and harvested after 3 days of infiltration. Proteins were extracted in buffer containing 125 mM Tris-HCl (pH 7.7), 0.25 mM EDTA, 2.5 mM MgCl_2_, 5% glycerol, 5 mM ATP, and protease inhibitors. YFP-HECT was then pulled down by using GFP-Trap agarose (ChromoTek). For purification of the proteasome, 4-week-old Arabidopsis plants were ground in extraction buffer and incubated overnight with anti-proteasome S2 antibody (Abcam) at 4°C. Protein complexes were then pulled down by using Protein A-agarose beads (Millipore). In vitro ubiquitination assays were performed by incubating the purified proteins (*i.e.* YFP-HECT or proteasomes) in 80 μl reaction buffer (125 mM Tris-HCl (pH 7.7), 0.25 mM EDTA, 2.5 mM MgCl_2_, 5 mM ATP, 1mM DTT, 10 μM NSC632836 deubiquitinase inhibitor) supplemented with 0.2 μg recombinant human E1 enzyme (BioVision), 0.2 μg recombinant E2 enzyme UbcH5c (Ubiquigent), and 10 μg recombinant human FLAG-ubiquitin (Boston Biochem) at 30°C for 18 h with shaking. To terminate the reaction, SDS sample buffer containing 50 mM DTT was added and incubated 80°C for 10 min before separating proteins by SDS-PAGE.

Protein-protein interaction assays

For detecting interactions between UPLs and NPR1, Agrobacterium carrying *35S:FLAG-UPL3*, *35S:MYC-UPL5*, or *35S:NPR1-GFP* constructs were collected and resuspended in infiltration buffer containing 10 mM MgCl_2_ and 10 μl/l 6-benzyladenine to OD_600_ = 0.3. *N. benthamiana* leaves were infiltrated and collected after 3 days of infiltration. For testing in vivo YFP-UPL3 and EIN3 interaction, Arabidopsis *35S:YFP-UPL3/upl3* and WT plants were treated with 100 μM MG132 for 2 h. Proteins were extracted as described above after which NPR1-GFP or YFP-UPL3 proteins were pulled down using GFP-Trap agarose (ChromoTek) according to the manufacturers’ instructions. Next, samples were heated at 70°C for 15 min in SDS sample buffer supplemented with 50 mM DTT before protein separation by SDS-PAGE and immunoblotting with anti-GFP and anti-EIN3 (*40*) antibodies.

For analysis of interaction between FLAG-EBF2 and YFP-UPL3, YFP-UPL3 was purified with GFP-Trap agarose from plants carrying *35S:YFP-UPL3*. Agarose beads were washed 3 times with wash buffer (10 mM Tris/Cl pH 7.5, 150 mM NaCl, 0.5 mM EDTA, protease inhibitors) before incubating in wash buffer containing cell-free synthesized FLAG-EBF2 (*41*) at 4°C for 1 h with rotation. Beads were washed extensively with wash buffer and then boiled for 10 minutes in SDS sample buffer containing 50 mM DTT. FLAG-EBF2 was detected by immunoblotting with an anti-FLAG-HRP antibody (Sigma-Aldrich).

For analysis of interaction between EIN3-GFP-FLAG and the proteasome, EIN3-GFP-FLAG was purified with GFP-Trap agarose from the indicated genotypes carrying *pEIN3:gEIN3-eGFP-3xFLAG*. Presence of the proteasome was detected using an S2 antibody (Abcam).

To test if interaction between UPL3 and EIN3 depends on EBF1/2, 10^6^ protoplasts from *35S:YFP-UPL3* (in *ein3*) or *35S:YFP-UPL3* (in *ebf1 ebf2 ein3*) plants were prepared and transformed with 100 μg of pEarleyGate 201/*35S:HA-EIN3* plasmid DNA as described previously (*42*). Next, proteins were extracted as described above, YFP-UPL3 protein purified using GFP-Trap agarose, and HA-EIN3 and YFP-UPL3 detected using anti-HA (ThermoFisher) and anti-GFP (Roche) antibodies, respectively.

Plant hormone analysis

SA content was determined according to a previously described method with specified modifications (*43*). In brief, fresh leaves were ground in liquid nitrogen and 0.1 g of sample was suspended in 4 ml of extraction buffer (1% (v/v) acetic acid in acetonitrile/water (4:1)) with stable isotope-labelled internal standards. Suspended samples were extracted, centrifuged and concentrated as described previously. Samples were purified by solid phase extraction using Oasis WAX cartridges (Waters Corp., Milford, MA, USA) from which SA was eluted with 3% (v/v) formic acid in acetonitrile. Following evaporation of each fraction, samples were analysed on an Agilent 1260–6410 Triple Quad LC/MS system (Agilent Technologies Inc., Santa Clara, CA, USA) equipped with a Capcell Pak ADME-HR S2 column (Osaka Soda Co. Ltd., Osaka, Japan).

Chromatin immunoprecipitation

Chromatin immunoprecipitation was performed as described previously but with minor modifications (*44*). A total of 500 mg tissue was crosslinked with 1% formaldehyde by vacuum infiltration for 15 mins at room temperature. Glycine was added to a final concentration of 125 mM to quench the crosslinking reaction and tissue was vacuum infiltrated for a further 5 mins. Crosslinked tissue was washed three times with ice-cold PBS before freezing in liquid nitrogen. For analyses nuclei were isolated and lysed as described (*44*), while sonication was performed using a BioRuptor Plus (Diagenode) for 10 cycles of 30 s ON, 30 s OFF at high power. NPR1-GFP and YFP-UPL3 were immunoprecipitated using an anti-GFP antibody (Abcam), EIN3 was immunoprecipitated using an anti-EIN3 antibody (*40*), and the proteasome with an anti-RPN10 antibody (Abcam). Enrichment at chromatin binding sites was analysed by qPCR using primers listed in Table S2.

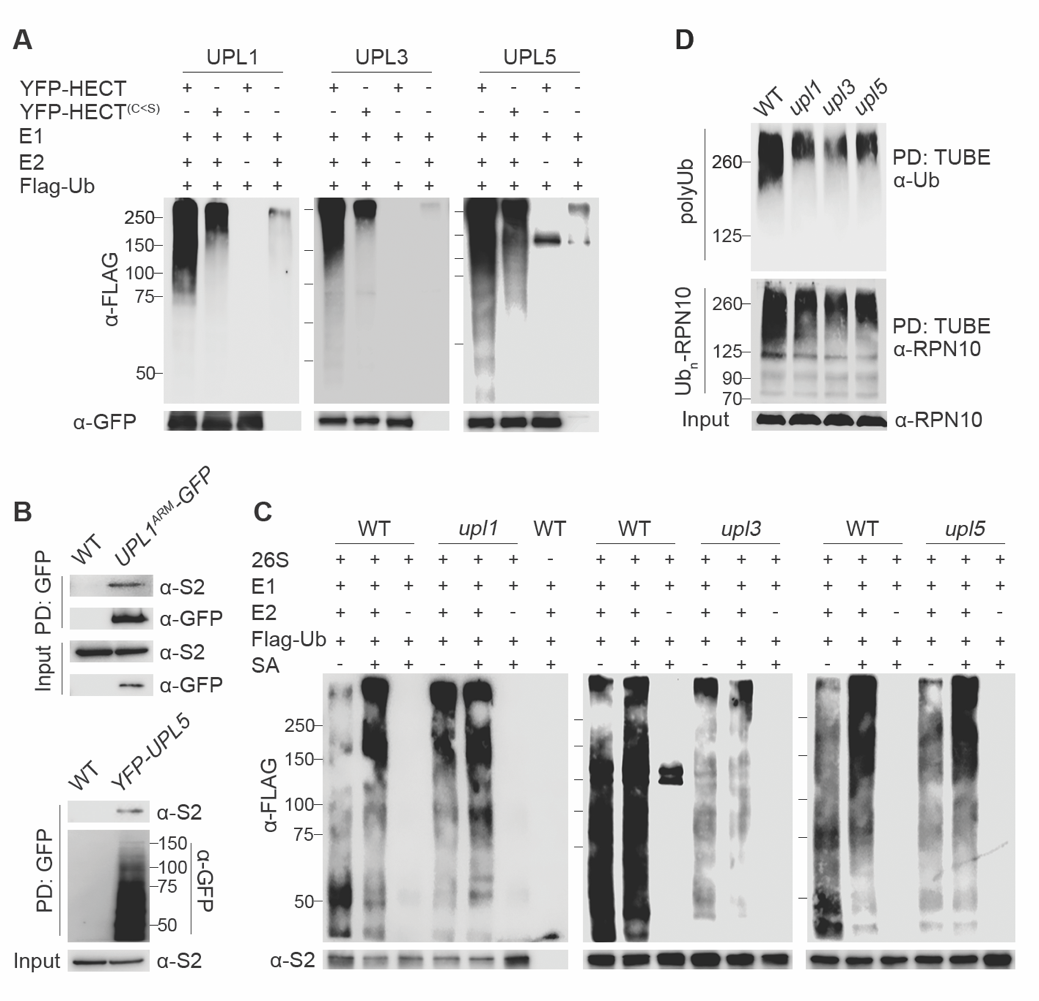

**Fig. S1. UPL ligases confer proteasome-associated ubiquitin ligase activity**

(**A**) UPLs exhibit ubiquitin ligase activity. *35S:YFP-HECT* constructs with or without mutation of the active-site cysteine were transiently expressed in tobacco leaves, while buffer-infiltrated leaves were used as a negative control. Purified proteins were incubated with E1 enzyme, E2 enzyme, FLAG-ubiquitin (FLAG-Ub) and ATP. FLAG-ubiquitin conjugates were detected by western blotting using an anti-FLAG antibody. Expression of YFP-HECT proteins were detected by using a GFP antibody. Note that HECT ligases utilise a two-step ubiquitination system that involves transfer of ubiquitin from the E2 enzyme to the HECT active site cysteine and finally to the substrate. Therefore, residual activity remained in HECT mutants as these proteins still interact with and facilitate activity of the E2 enzyme. (**B**) UPL1 and UPL5 physically interact with the proteasome *in planta*. UPL1^ARM^-GFP and YFP-UPL5 proteins were immunoprecipitated from seedlings expressing *35S:UPL1^ARM^-GFP* and *35S:YFP-UPL5*, respectively, whereas WT plants were used as a negative control. Immunoprecipitated proteins were analysed by western blotting using antibodies against S2 (*i.e.* proteasome regulatory non-ATPase subunit RPN1) and GFP. (**C**) UPL3 predominantly contributes to proteasome-associated ubiquitin ligase activity. Adult plants were treated with (+) or without (-) 0.5 mM SA for 24 hours. Proteasomes were immunoprecipitated with an anti-S2 antibody, whereas pull downs in absence of antibodies were performed as a control. Ubiquitination assay was performed and analysed as in (A). (**D**) UPLs contribute substantially to global cellular levels of ubiquitin conjugates. Seedlings were treated with 0.5 mM SA and 0.1 mM MG132 for 6 hours. Ubiquitinated substrates were pulled down using His-TUBE and analysed by immunoblotting with a ubiquitin antibody. Polyubiquitinated and input RPN10 are indicated.

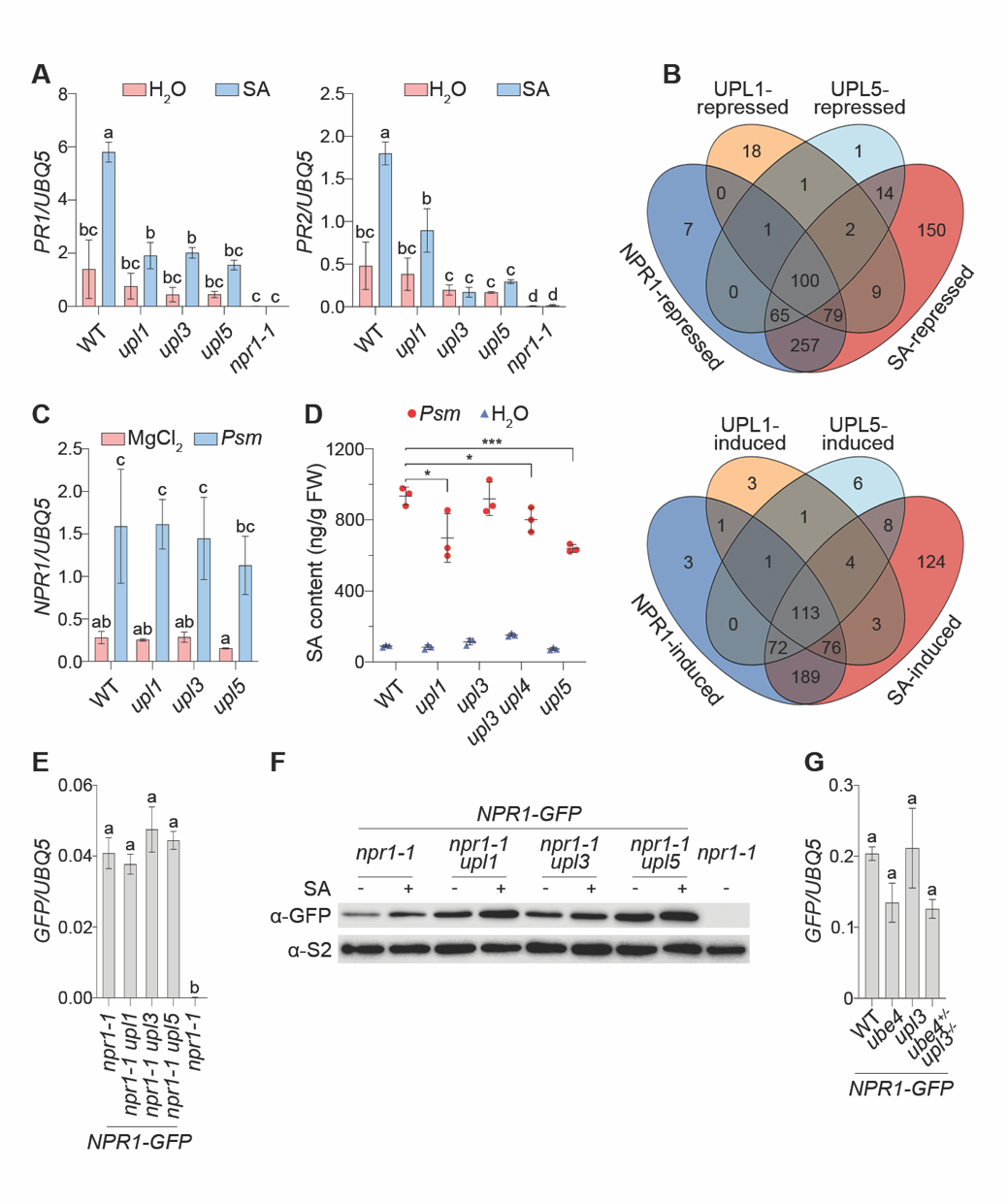

**Fig. S2. UPLs regulate SA- and NPR1-dependent signalling**

**(A**) Mutant *upl* plants exhibit reduced *PR* gene expression. Adult plants were treated with 0.5 mM SA or H_2_O for 24 hours. Expression of *PR* genes was normalised to constitutively expressed *UBQ5*. Data represent mean ± SD, lowercase letters indicate statistically significant differences between samples (Tukey HSD ANOVA test; α = 0.05, n = 3). (**B**) Venn diagram of SA-responsive genes differentially expressed in WT versus the *upl1*, *upl5*, and *npr1* mutants. Overlap of SA-responsive (in WT), UPL1-, UPL5-, and NPR1-regulated genes is indicated. (**C**) *NPR1* mRNA levels are normal in the *upl* mutants. Leaves were inoculated with 10^6^ cfu/ml *Psm* ES4326 or 10 mM MgCl_2_, samples were collected at 48 hours after infection. Expression of *NPR1* was normalised to *UBQ5*. Data represent mean ± SD, lowercase letters indicate statistically significant differences between samples (Tukey HSD ANOVA test; α = 0.05, n = 3). (**D**) SA levels are reduced in *upl* mutants. Leaves were infected and collected as in (C). Data represent mean ± SD, asterisk indicate statistically significant differences between samples (two-tailed t test, **p* ≤ 0.05, ****p* ≤ 0.001, n = 3). (**E**) Equal *NPR1-GFP* mRNA accumulation in the *upl* mutants. Expression of *NPR1-GFP* in the indicated genotypes was normalised to *UBQ5*. Data represent mean ± SD, lowercase letters indicate statistically significant differences between samples (Tukey HSD ANOVA test; α = 0.05, n = 3). (**F**) Constitutively expressed *NPR1-GFP* without UTRs accumulates in the *upl* mutants. Seedlings were treated with 0.5 mM SA or H_2_O for 6 hours. The *npr1-1* mutant was used as a negative control. NPR1-GFP protein was detected by western blotting, whereas protein levels of S2 were used as a loading control. (**G**) Equal *NPR1-GFP* mRNA accumulation in *ube4* and *upl3* single and double mutants. Expression of *NPR1-GFP* in the indicated genotypes was analysed as in (E).

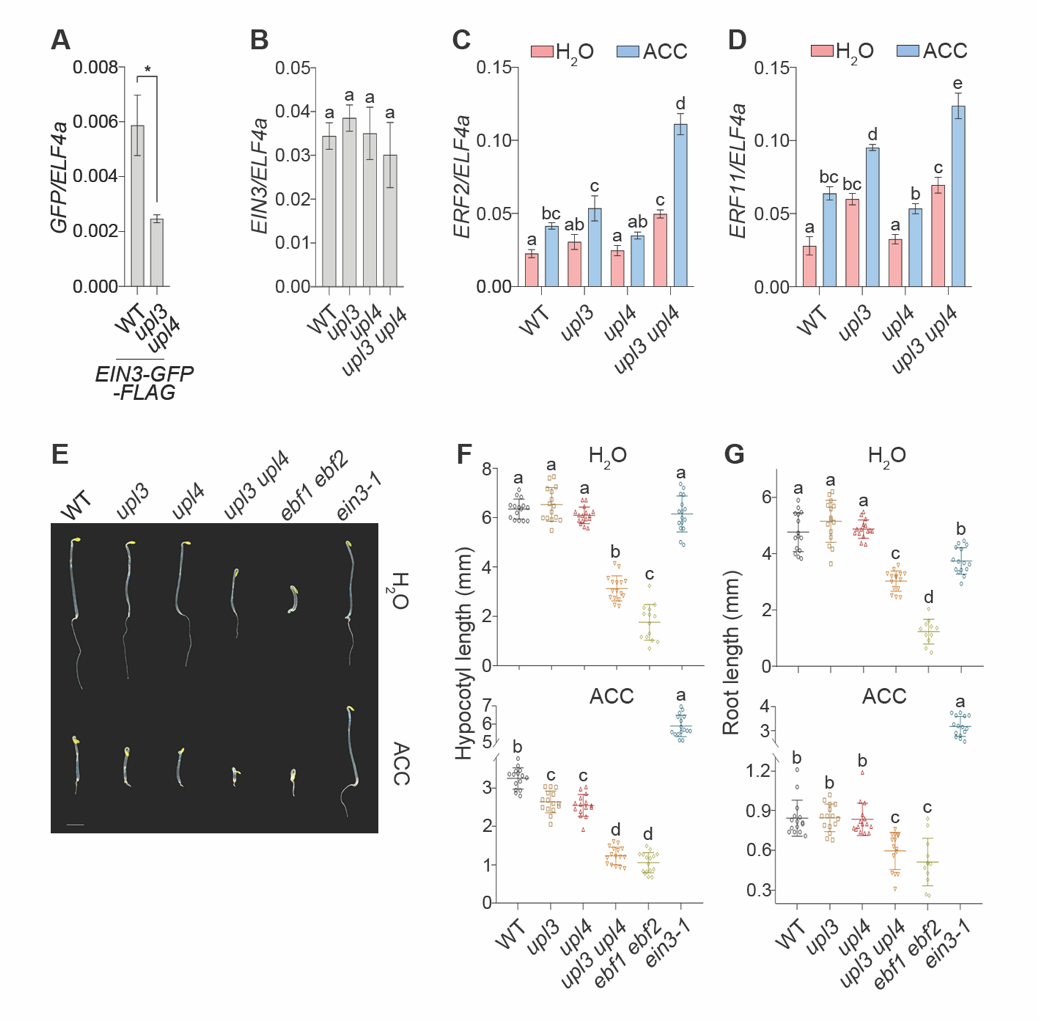

**Fig. S3. UPL3/4 exhibit constitutive ET responses**

(**A**) Accumulation of *EIN3-eGFP-3xFLAG* mRNA in WT and *upl3 upl4*. Expression of GFP in the indicated genotypes was normalized to *ELF4a*. Data represent mean ± SD, asterisk (two-tailed t test, *p ≤ 0.05, n = 3) indicate statistically significant between samples. (**B**) Equal *EIN3* mRNA accumulation in WT and *upl* mutants. Expression of *EIN3* was normalised to *ELF4a*. Data represent mean ± SD, lowercase letter indicates no statistically significant differences between samples (Tukey HSD ANOVA test; α = 0.05, n = 3). (**C** and **D**) EIN3 target genes are up-regulated in the *upl3 upl4* mutant. Seedlings of the indicated genotypes were treated with 50 μM ACC or H_2_O for 3 hours. Expression of *ERF2* (C) and *ERF11* (D) were normalised to *ELF4a*. Data represent mean ± SD, lowercase letters indicate statistically significant differences between samples (Tukey HSD ANOVA test; α = 0.05, n = 3). (**E** to **G**) Mutant *upl3 upl4* plants display constitutive ET response phenotypes. Etiolated Seedlings were grown on MS medium supplemented with or without 10 μM ACC. Hypocotyl (F) and root (G) lengths of 4-day-old seedlings were measured. Data represent mean ± SD, lowercase letters indicate statistically significant differences between samples (Tukey HSD ANOVA test; α = 0.05, n = 15)

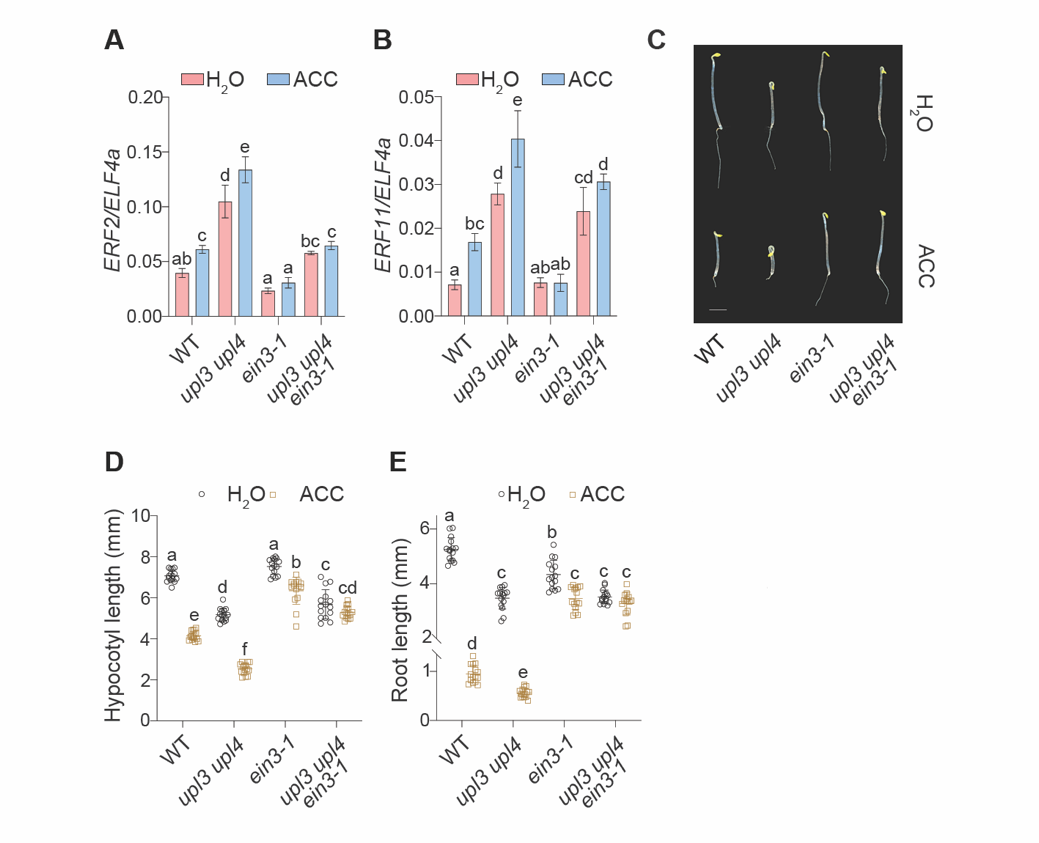

**Fig. S4 ET responses of *upl3/4* mutants are dependent on *EIN3*.**

(**A** and **B**) Mutation of *EIN3* in the *upl3 upl4* mutant suppressed elevated expression levels of *ERF*s. Expression of *ERF2* (A) and *ERF11* (B) was normalised to *ELF4a*. Data represent mean ± SD, lowercase letters indicate statistically significant differences between samples (Tukey HSD ANOVA test; α = 0.05, n = 3). (**C** to **E**) ET-responsive phenotypes of 4-day-old seedlings of the indicated genotypes were analysed. Hypocotyl (D) and root (E) lengths of 4-day-old seedlings were measured. Data represent mean ± SD, lowercase letters indicate statistically significant differences between samples (Tukey HSD ANOVA test; α = 0.05, n = 15)

**Table S1. Key resources**

| Reagent type or resource  (species) or resource | Designation | Source | Identifier |
| --- | --- | --- | --- |
| Antibody | Mouse monoclonal anti-GFP | Roche | Cat# 11814460001 |
| Antibody | Rabbit polyclonal anti-NPR1 | (*34*) | N/A |
| Antibody | Rabbit polyclonal anti-GFP (ChIP grade) | Abcam | Cat# ab290 |
| Antibody | Mouse monoclonal anti-Ubiquitin (FK2) | Millipore | Cat# 04–263 |
| Antibody | Mouse monoclonal anti-FLAG M2-Peroxidase (HRP) | Sigma-Aldrich | Cat# A8592 |
| Antibody | Rabbit polyclonal anti-Proteasome 26S S2/PSMD2 | Abcam | Cat# Ab98865 |
| Antibody | Rabbit polyclonal anti-Proteasome 19S S5A/ASF | Abcam | Cat# Ab56851 |
| Antibody | Mouse monoclonal anti-Myc | Sigma-Aldrich | Cat# M5546 |
| Antibody | Rabbit polyclonal anti-EIN3 | (*40*) | N/A |
| Antibody | Mouse monoclonal anti-HA | ThermoFisher | Cat# 26183 |
| Recombinant DNA | pENTR-D-TOPO | Invitrogen | Cat# K240020 |
| Recombinant DNA | pEarleyGate 103 | (*45*) | Cat# CD3-685 |
| Recombinant DNA | pEarleyGate 104 | (*45*) | Cat# CD3-686 |
| Recombinant DNA | pEarleyGate 201 | (*45*) | Cat# CD3-687 |
| Recombinant DNA | pEarleyGate 202 | (*45*) | Cat# CD3-688 |
| Recombinant DNA | pEarleyGate 203 | (*45*) | Cat# CD3-689 |
| Recombinant DNA | pK7FWG2 | (*46*) | N/A |
| Recombinant DNA | pCAB1:YFP-GW | (*36*) | CD3-1938 |
| Arabidopsis thaliana | *upl1* | (*33*) | SALK_063972 |
| Arabidopsis thaliana | *upl3* | (*33*) | SALK_ 035524 |
| Arabidopsis thaliana | *upl4* | (*33*) | SALK_040984 |
| Arabidopsis thaliana | *upl5* | (*33*) | SALK_116446 |
| Arabidopsis thaliana | *35S:YFP-UPL3* | (*33*) | N/A |
| Arabidopsis thaliana | *npr1-0* | (*34*) | SALK_204100 |
| Arabidopsis thaliana | *ube4-2* | (*47*) | SAIL_713_A12 |
| Arabidopsis thaliana | *npr1-1* | (*35*) | N/A |
| Arabidopsis thaliana | *ein3-1* | (*32*) | N/A |
| Arabidopsis thaliana | *ebf1-1 ebf2-1* | (*32*) | N/A |
| Commercial assay or kit | SuperScript II | Invitrogen | Cat# 18064014 |
| Commercial assay or kit | QuikChange Site-Directed Mutagenesis Kit | Agilent | Cat# 200519 |
| Commercial assay or kit | GFP-Trap A | Chromotek | Cat# gta-20 |
| Chemical compound, drug | MG132 | Cayman Chemical | Cat# 10012628 |
| Chemical compound, drug | Cycloheximide | Sigma-Aldrich | Cat# C7698 |
| Chemical compound, drug | Sodium salicylate | Sigma-Aldrich | Cat# S3007 |
| Chemical compound, drug | 1-Aminocyclopropanecarboxylic acid | Sigma-Aldrich | Cat# A3903 |
| Software, algorithm | Strand NGS | Avadis | N/A |

Table S2. List of primers used in this study

| Name | Purpose | Sequences (5’-3’) | Source |
| --- | --- | --- | --- |
| *PR1* | qPCR | F: CTAAGGGTTCACAACCAGGC  R: AAGGCCCACCAGAGTGTATG | (*34*) |
| *PR2* | qPCR | F: CAGATTCCGGTACATCAACG  R: AGTGGTGGTGTCAGTGGCTA | (*34*) |
| *NPR1* | qPCR | F: CTAAAACCGTGGAACTCGGG  R: TCTCTTGTATTTCCATGTACCTTTGCT | (*34*) |
| *UBQ5* | qPCR | F: CCAAGCCGAAGAAGATCAAG  R: ACTCCTTCCTCAAACGCTGA |  |
| *UPL3* | qPCR | F: CGTCTTCTTGTGCTGCTGTT  R: GAGCACAGCCATAAGAGCAC |  |
| *UPL4* | qPCR | F: TCGCCCGTTCTTGTCCTATT  R: GTTCATGCCCTCCAGACTCT |  |
| *ERF1* | qPCR | F: GAGGAAACACTCGATGAGACG  R: GGAGCGGTGATCAAAGTCAC | (*48*) |
| *ERF2* | qPCR | F: CGAGCCAACTGAGAACTTTA  R: CAAACCCTAGCTCCATTCTT |  |
| *ERF11* | qPCR | F: GTACTTTCGACACTCCTGAAG  R: CAGATCCGAGGTTGAGATTAG |  |
| *ELF4a* | qPCR | F: TCATAGATCTGGTCCTTGAAAC  R: GGCAGTCTCTTCGTGCTGAC | (*48*) |
| *EIN3* | qPCR | F: GGATAAGGGTAAAGAAGGTGTT  R: GACCATTACGATCAAACCTAAC |  |
| *GFP* | qPCR | F: AAGCTGACCCTGAAGTTCATCTGC  R: CTTGTAGTTGCCGTCGTCCTTGAA |  |
| *EBF2* | Cell-free protein synthesis | F: CCAGCAGGGAGGTACTATGTCTGGAATCTTCAGA  R:  CCTTATGGCCGGATCCAAGAGCTCTTTTTTTTTTTTAGTAGAGTATATCGCA |  |
| *PR1* promoter *as1* element | ChIP qPCR | F: AGTGTATACAATGTCAATCGGTGATCTT  R: GCCGCCACATCTATGACGTA | (*34*) |
| *ERF1* promoter | ChIP qPCR | F: GGGGGCATGTATCTTGAATC  R: TGCTGGATCAACTCAACAAAA | (*49*) |
| UPL1^HECT^ | Cloning | F: CACCATGCATGGTTCCTCATCTAAAAC  R: TCAAGCAAACCCAAAACCTTC |  |
| UPL1^HECT (C<S)^ | Cloning | F: CTGCCATCAGCTCATACATCTTTTAACCAACTAGACCTC  R: GAGGTCTAGTTGGTTAAAAGATGTATGAGCTGATGGCAG |  |
| UPL3^HECT^ | Cloning | F: CACCATGTTGTGCAGTGGAAGTCTTC  R: TTATGAGAGGTCGAACGATCC |  |
| UPL3^HECT (C<S)^ | Cloning | F: TAAGGTAGTTTGCGGAAGTCATGACACTGGGCAA  R: TTGCCCAGTGTCATGACTTCCGCAAACTACCTTA |  |
| UPL5^HECT^ | Cloning | F: CACCATGATTTATCAAGGTGCTAAGGGAC  R: TCACCATTTACCGAAACTGGAGCTG |  |
| UPL5^HECT (C<S)^ | Cloning | F: GTATACAGAGACGGTAAAAGGAAGTATGAGACAATGGAAGA  R: TCTTCCATTGTCTCATACTTCCTTTTACCGTCTCTGTATAC |  |

42.	L. L. Hansen, G. van Ooijen, Rapid Analysis of Circadian Phenotypes in Arabidopsis Protoplasts Transfected with a Luminescent Clock Reporter. *J Vis Exp*, (2016).
